# High prevalence of the antibody against Syncytin-1 in schizophrenia

**DOI:** 10.1101/266049

**Authors:** Yurina Hibi, Kaori Asamitsu, Hiroyo Matsumura, Takaomi Sanda, Yuki Nakahira, Kenji Arimoto, Shinsuke Nakanishi, Kazunori Maekawa, Tatsuo Akechi, Noriyuki Matsukawa, Terutaka Fukaya, Yutaka Tomita, Nariaki Iijima, Hiroyuki Kato, Takashi Okamoto

## Abstract

Both genetic and environmental factors have been considered causative agents for schizophrenia (SZ). However, no single gene has been shown responsible for the development of SZ. Furthermore, the pathophysiological roles of environmental factors including psychological stress, autoimmunity, and microbial infection have not been fully understood. Previous studies have suggested the involvement of one of the human endogenous retroviruses (HERVs), HERV-W, in SZ. In this study, prevalence of antibodies against the HERV-W Syncytin-1 protein was examined using a newly developed ELISA test. Fifty percent of patients with SZ (24 out of 48 cases) were antibody-positive, with a specificity of greater than 95% (less than 5% of control cases, 3 out of 79). No significant effect of medication was evident, nor did any SZ cases become seropositive after diagnosis. These findings indicate a possible involvement of HERV-W expression in the development of SZ and support its applicability to laboratory diagnoses.

## Introduction

Schizophrenia (SZ) is a major psychiatric disorder that affects 0.5 to 1.0% of the populations in most countries, irrespective of socio-economic conditions (Owen *et al*, 2016). Diagnosis of SZ is often made by experienced psychiatric experts in accordance with the Diagnostic and Statistical Manual of Mental Disorders, 5^th^ edition (DSM-5) (American Psychiatric Association, 2013) or the International Classification of Diseases, 10^th^ edition (ICD-10) (WHO, 1992); neither includes objective diagnostic measures, such as laboratory tests. In addition, differential diagnosis of closely related psychiatric conditions such as bipolar disorder (BD) and schizoaffective disorder is not currently clear. Thus, non-invasive blood testing is needed to facilitate early and objective diagnosis of psychopathic patients.

Although a number of genome-wide association studies (GWASs) have demonstrated a genetic predisposition to SZ (Lichtenstein *et al*, 2009; Purcell *et al*, 2014; Takata *et al*, 2017), no single gene has been shown to be responsible. Other studies have revealed complex interactions between the nervous and immune-inflammatory systems (Khandaker *et al*, 2015; Fillman *et al*, 2013). Interestingly, a recent epidemiological study confirmed a link between SZ and autoimmunity (Heath *et al*, 1967; Chen *et al*, 2012; Benros *et al*, 2014; Cremaschi *et al*, 2017). For instance, a nationwide risk-assessment study revealed a cross relationship between these two pathologies (Benros *et al*, 2014; Cremaschi *et al*, 2017) with an increased risk of subsequent autoimmune diseases in individuals with SZ. In addition, subsequent studies indicated a overlap in common SNPs between schizophrenia and autoimmune diseases (Hoeffding *et al*, 2017). These findings suggest the presence of a common etiological mechanism.

The human genome contains a number of retrovirus-related sequences, including human endogenous retroviruses (HERV), long interspersed nuclear elements (LINE) and short interspersed nuclear elements (SINE), that comprise nearly 45% of the entire genome (Kazazia *et al*, 2017). Most of these sequences are considered to have been derived from previously infected ecotropic retroviruses even before the divergence of *Homo sapiens* from the rest of the mammals. Conservation of these sequences over time may be indicative of their pivotal roles in cell differentiation or immunological and neuronal diversification (Kazazia *et al*, 2017). HERV sequences generally degenerate and lose open reading frames throughout molecular evolution (Kazazia *et al*, 2017). However, one such HERV, HERV-W, contains a conserved long open reading frame of 538 amino acids (AAs), named Syncytin-1 (Mi *et al*, 2000; Lavialle *et al*, 2013). The expression of *syncytin-1* is considered essential for placenta organogenesis as its expression has been demonstrated to induce the syncytiotrophoblast (Mi *et al*, 2000; Lavialle *et al*, 2013) and is tightly regulated by an epigenetic mechanism (Matousková *et al*, 2016). Gene expression of HERV-W is aberrantly induced under extreme conditions such as viral infection and carcinogenesis (Kassiotis, 2014; Nellåker *et al*, 2006), playing as a molecular marker for the cell damage and its transformation. Thus, presence of HERV is considered beneficial for the maintenance of species.

HERV-W was first isolated from the cerebrospinal fluid of patients with multiple sclerosis and was thus initially named the multiple sclerosis-associated retrovirus (MSRV) (Kurth, 1986). It is possible aberrant expression of HERV-W might trigger the aberrant immune responses followed by chronic inflammation and the neurodegenerative changes characteristic for multiple sclerosis (MS). Furthermore, HERV-W is considered to be involved in a number of other diseases. Interestingly, expression of the HERV-W sequence has been previously observed in the brain tissue of SZ patients as well as healthy brain tissues (Yolken *et al*, 2000; Karlsson *et al*, 2001; Frank *et al*, 2005). In addition, expression of HERV-W sequences were detected in the cerebrospinal fluid of SZ patients (Frank *et al*, 2005). More specifically, the Syncytin-1 protein and its mRNA were detected in the sera of SZ patients (Perron *et al*, 2008; Perron *et al*, 2012). However, no evidence for the presence of the anti-Syncytin-1 antibody in these patients has yet to be demonstrated. In this study, we demonstrate for the first time that antibodies against the Syncytin-1 protein are present in the blood of individuals with psychiatric diseases. Implication of the presence of Syncytin-1 autoantibody will be discussed.

## Results

We have examined the prevalence of anti-Syncytin-1 antibody among neuropsychiatric patients including SZ [n=48], bipolar disorder (BD) [n=18], MS [n=17] and control subjects [n=79]. There were no significant differences in the male to female ratio or average age among different disease categories (Table 1). The anti-Syncytin-1 antibody was also examined in BD known crossly related to SZ (Owen *et al*, 2016; American Psychiatric Association, 2013; WHO, 1992). To initially screen the anti-Syncytin-1 antibody, we applied Western blotting using semi-purified recombinant full-length Syncytin-1 protein (Appendix Fig. S1). However, because of the instability of semi-purified proteins due to high hydrophobicity upon Western blotting (e.g., the broadening of the immune-reacted bands) as well as the technical difficulty in purifying the protein, we attempted to develop a novel antibody screening method using ELISA and antigenic peptides derived from Syncytin-1.

**Table 1.**
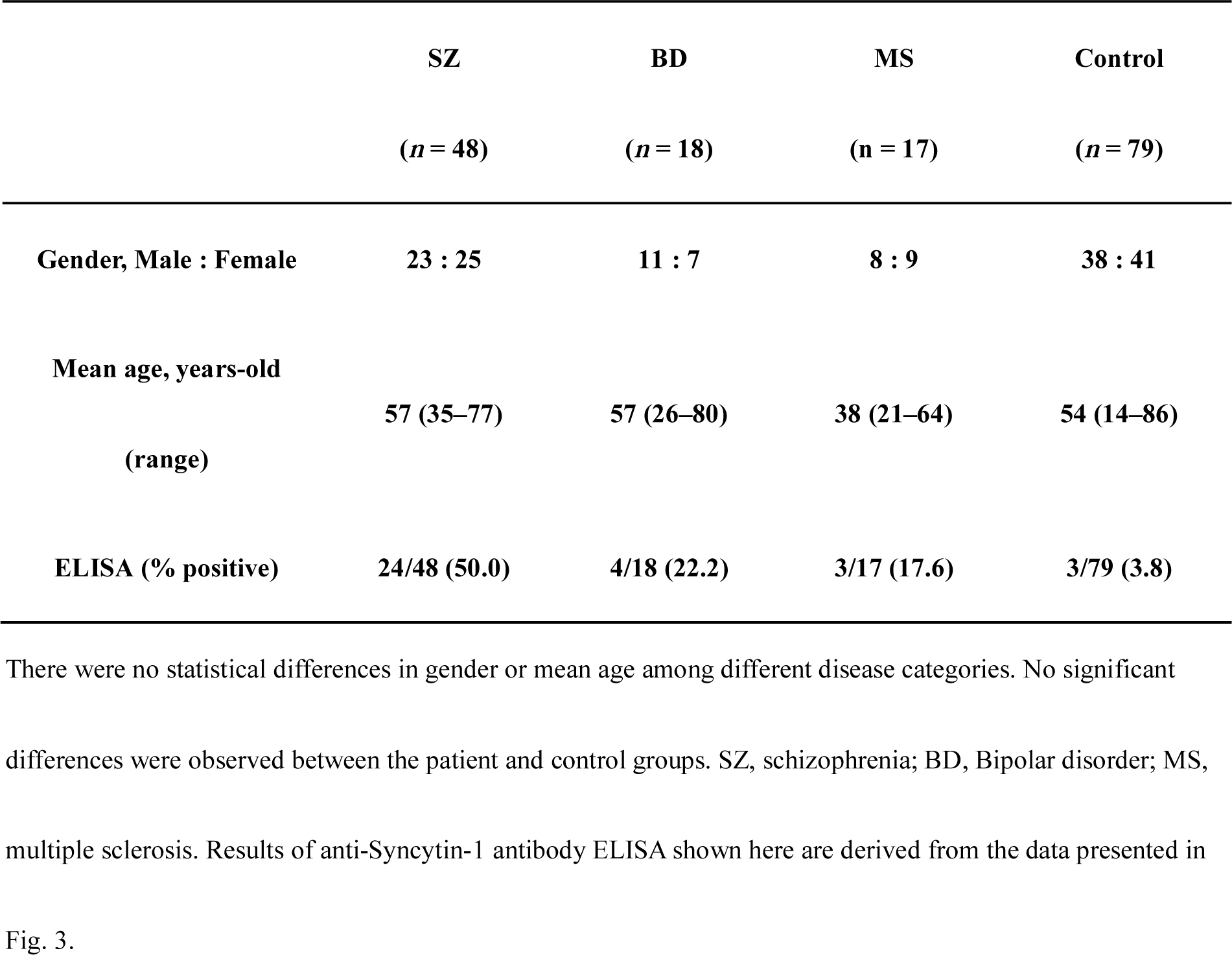
Prevalence of anti-Syncytin-1 antibody in schizophrenia, bipolar disorder, and multiple sclerosis.

In order to develop a specific ELISA system for anti-Syncytin-1 antibody detection, we determined commonly-recognized Syncytin-1 epitopes using a Syncytin-1 peptide array (Fig. 1). The results indicated that most of the positive sera reacted to the peptides containing various epitope regions (Fig. 1a). The shaded areas in Fig. 1B indicated the Syncytin-1 peptide regions that were highly immune-reactive to each patient’s serum (quantitation of the spots demonstrated in Fig. 1a). Each peptide sequence is listed in Appendix Table S2. Among the 48 SZ patients there was no significant difference in the presence of pathognomonic symptoms such as hallucinations, delusions, thought disorders, movement disorders, “flat affect” or reduced speaking, irrespective of the presence of anti-Syncytin-1 antibody (Table 1).

**Figure 1.**
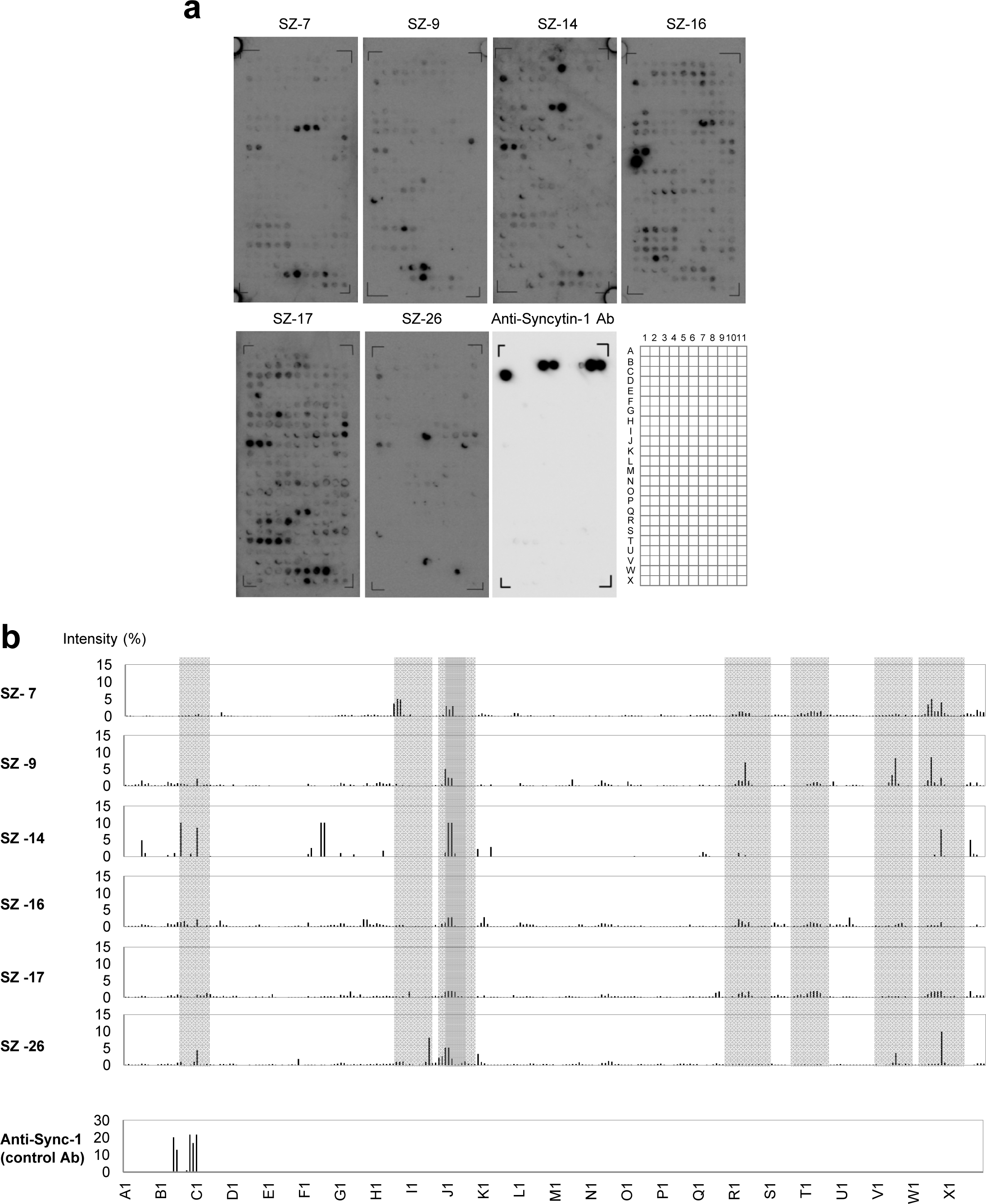
Determination of antigenic epitopes of Syncytin-1 using sera from 6 schizophrenia patients. (a) Immunoblot of Syncytin-1 peptide array for 6 seropositive schizophrenia (SZ) cases. In the array, each peptide spot denotes a 12 amino acids (AA)-long peptide with a frameshift of 2 AA throughout the entire 538 AA peptide (total 264 spots). The positive control of anti-Syncytin-1 antibody is included. (b) Quantitative representation of serum reactivity to each Syncytin-1 peptide. The data were obtained from the results in Fig. 1a. The common immune-reactive regions are shown as shaded areas.

Based on these serological findings, we have determined the antigenic peptide for ELISA screen (Fig. 2). Among the eight peptides of Syncytin-1, only the peptide #8 exhibited anti-Syncytin-1 antigenicity, and the inclusion of other peptides unexpectedly exhibited interfering effects (Fig. 2b). The peptide competition experiments in Fig. 2c show that the immune-reactivity of most positive sera against Syncytin-1 peptide #8 was inhibited only by the homologous peptide and not by the other peptides. We do not currently know the reason for the preferred antigenicity in peptide #8. However, since this peptide region is located in the cytoplasm (Mi *et al*, 2000; Lavialle *et al*, 2013) and is unlikely to be exposed, the antibody response could not have been elicited unless cells were destroyed to expose the membrane protein fractions including Syncytin-1.

**Figure 2.**
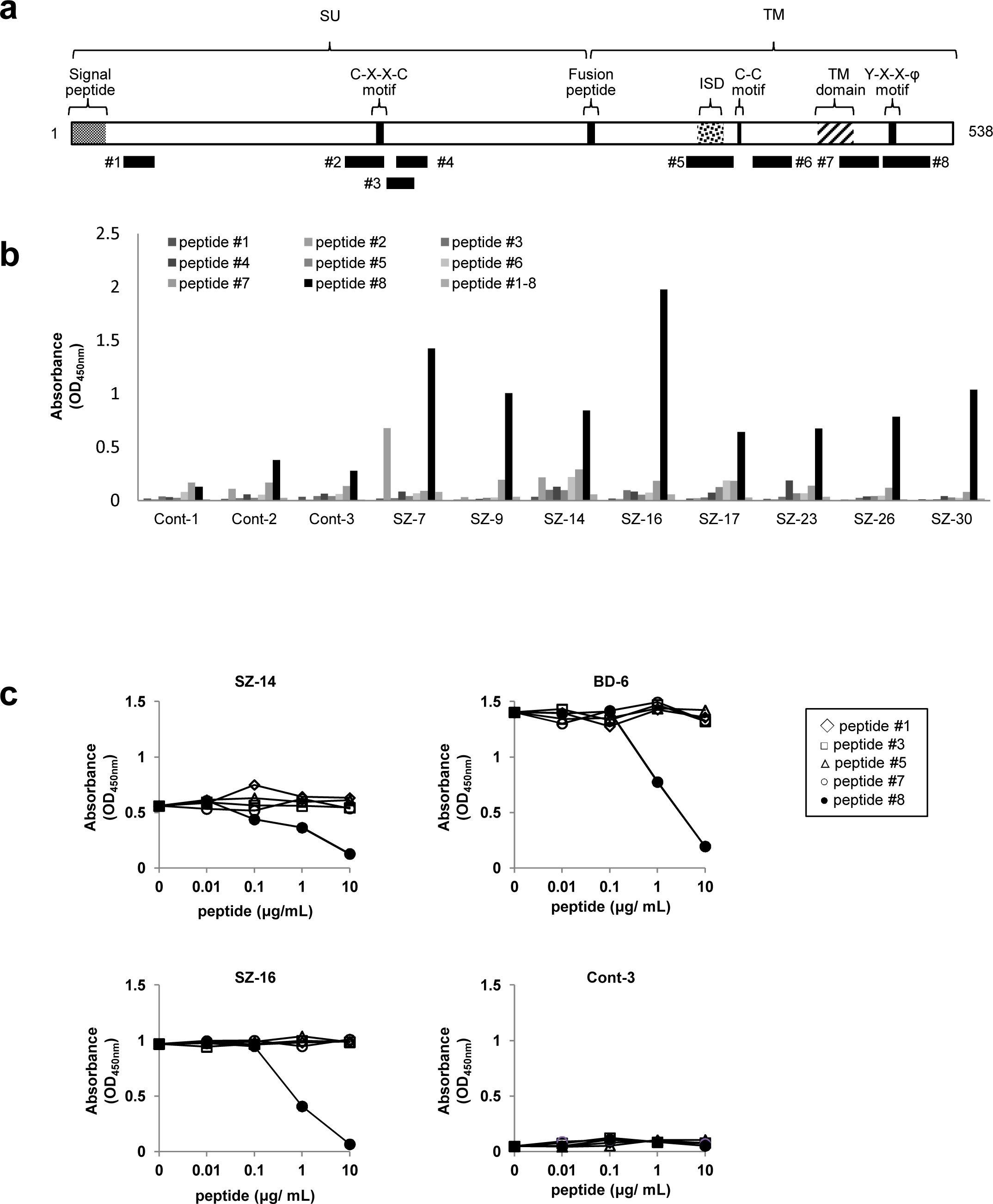
Establishment of ELISA system detecting anti-Syncytin-1 antibody among psychiatric cases. (a) The primary structure of HERV-W Syncytin-1 protein and locations of putative functional domains. Signal peptide (1–20 AA), C-X-X-C motif (186–189 AA), fusion peptide (314–317 AA), immunosuppressive domain (ISD; 380–396 AA), C-C motif (404–405 AA), TM domain (455–477 AA), Y-X-X-φ motif (498–501 AA). SU, surface area; TM, transmembrane region. The eight common immunogenic epitope regions are depicted as solid bars: #1, 30–58 AA; #2, 165–188 AA; #3, 192–208 AA; #4, 198–216 AA; #5, 375–403 AA; #6, 415–438 AA; #7, 467–490 AA; and #8, 493–521 AA. (b) ELISA reactivity of sera from SZ patients. The immune-dominant epitopes recognized by the anti-Syncytin-1 antibody were quantified by ELISA with the 8 peptides depicted in Fig. 2A. Note that all the anti-Syncytin-1 antibody-positive sera demonstrated here recognized the peptide #8 (Syn493–521) located in the C-terminal region of the Syncytin-1 protein. (c) Results of peptide competition ELISA. In order to assess the specificity of anti-Syncytin-1 ELISA detection system using Syncytin-1 peptide #8, two SZ sera (SZ-14 and SZ-16), one BD serum (BD-6), and a control serum (Cont-3) were examined. They were pre-incubated with various peptides at different concentrations before adding to the ELISA plate containing the #8 peptide. Note that significant competition was detected only with the homologous peptide.

Fig. 3a demonstrates the results of anti-Syncytin-1 antibody using ELISA system with sera from SZ, BD, MS, and controls. The antibody cut-off value was determined by the ROC curve (Fig. 3b). As summarized in Table 1 (bottom line), the anti-Syncytin-1 seroprevalence in SZ, BD and MS was 50.0% (24/48), 22.3% (4/18) and 17.6% (3/17), respectively; whereas that in controls was 3.8%. All positive sera against Syncytin-1 in Western blotting were also positive in ELISA. Some samples exhibiting the negative result in Western blotting showed positive reactivity in ELISA, indicating the higher sensitivity for ELISA. We thus used the ELISA system to examine the anti-Syncytin-1 antibody in 9 schizophrenic patients over time and found that the anti-Syncytin-1 antibody reactivity was largely stable except for a few cases, in which the immunoreactivity varied slightly in accordance with physical or mental stress (Fig. 4). Since HDAC inhibitors, such as valproic acid (VPA), are known to inhibit the maintenance of histone deacetylation and could possibly induce expression of repressed genes such as HERVs (Matousková *et al*, 2016; Diem *et al*, 2012), we have assessed the effects of VPA administration on the anti-Syncytin-1 antibody in SZ patients. However, there was no significant difference in antibody prevalence, irrespective of the VPA treatment of psychiatric patients. There was no subject who exhibited seropositivity in 9 patients under the VPA treatment to prevent seizures after brain surgery (data not shown).

**Figure 3.**
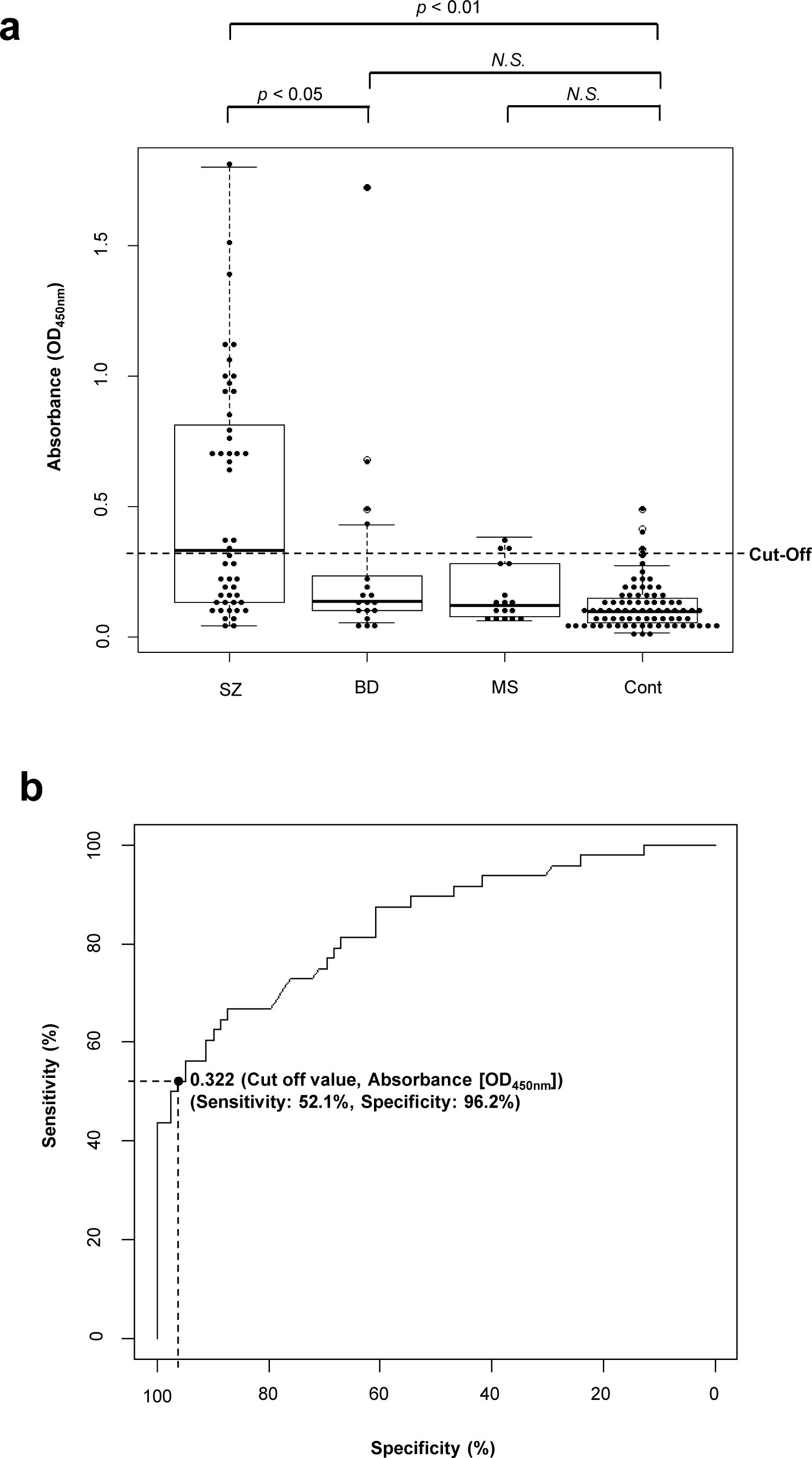
High prevalence of anti-Syncytin-1 antibody in schizophrenia. (a) ELISA plots of anti-Syncytin-1 antibody for SZ, BD, MS, and control cases. Note that SZ exhibited significantly higher antibody prevalence than BD, MS, and control cases. Antibody positivity was determined by the optical density (OD) value for each serum, with a cut-off value of 0.322 OD at 450 nm, as determined by ROC analysis. The seroprevalence values for each disease category are summarized in Table 1. Statistical significance was assessed by Wilcoxon’s rank sum test. The median values are indicated by the central horizontal line, and the data distribution is diagrammatically represented as a box-and-whisker plot. (b) ROC curve for detection of anti-Syncytin-1 antibody. The absorbance data at OD 450 nm from 48 SZ and 79 control sera were utilized to depict ROC curve. The cut-off value of 0.322 distinguishes positive and negative samples with a sensitivity of approximately 52% and a specificity of 96%.

## Discussion

Although SZ affects a large proportion of the general population throughout the world, its diagnosis has usually been made only by psychiatric experts, and most patients do not initially have direct access to medical care (Owen *et al*, 2016). Thus, novel diagnostic measures are needed to detect SZ upon initial manifestation of symptoms for further medical care of such patients. Our present findings of anti-Syncytin-1 antibody detection in psychiatric patients would meet such a need. Further verifications with extended study subjects should verify the significance of anti-Syncytin-1 antibody detection. These include determining whether this antibody is (i) produced as a result of various therapeutic measures or medical interactions that could modify the antibody production, (ii) detected in younger SZ patients upon its initial diagnosis, (iii) detected in other conditions related to SZ such as BD and schizoaffective disorders, (iv) produced as cause/effect of such pathologic conditions or (v) detected in SZ patients in other countries. As far as this initial study is concerned, there was no psychiatric case in which the antibody was detected after the diagnosis of SZ or BD was made. We observed subtle fluctuations of the antibody titer during the disease course (Fig. 4), which needs to be further verified. Regarding the influence of epigenetic modifiers such as VPA for expression of HERV-W (Diem *et al*, 2012) and production of anti-Syncytin-1 antibody, we did not observe a significant correlation between the use of VPA and the prevalence of anti-Syncytin-1 antibody (data not shown).

**Figure 4.**
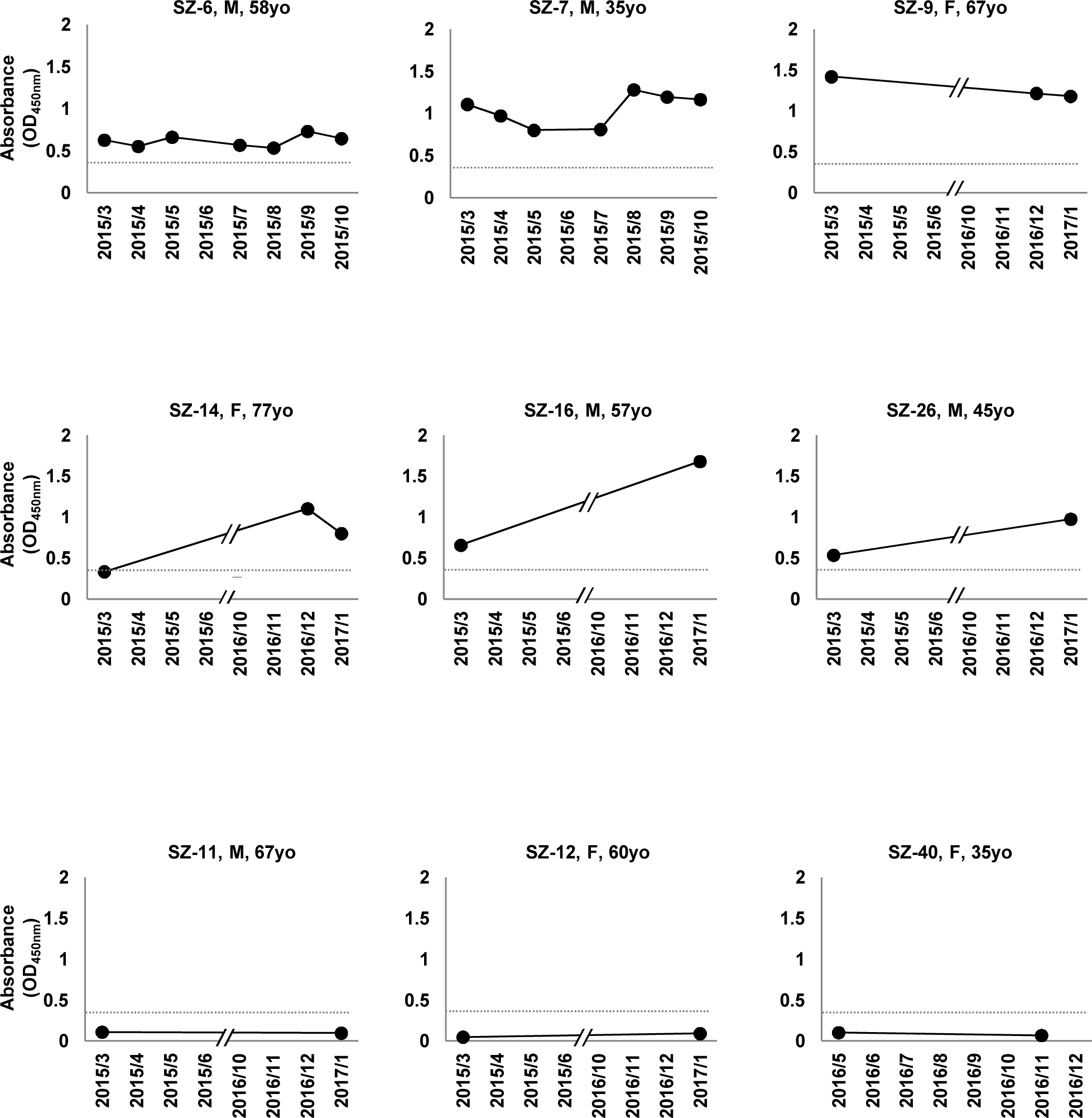
Temporal changes of the anti-Syncytin-1 antibody reactivity in schizophrenia patients. The serum anti-Syncytin-1 antibody titer was quantified by the ELISA system using the samples from 9 SZ patients during the disease course.

Previous reports suggested a possible relationship between autoimmunity and SZ (Heath *et al*, 1967; Chen *et al*, 2012). In addition to the recent epidemiological studies (Benros *et al*, 2014; Cremaschi *et al*, 2017), a number of studies reported the presence of autoantibodies such as antinuclear antibodies and anti-*N*-methyl-D-aspartic acid receptor (NMDAR) antibodies in SZ patients (van Mierlo *et al*, 2015; Kristiansen *et al*, 2007; Coyle, 2006). On the other hand, patients with paraneoplastic syndrome associated with ovarian teratome exhibiting anti-NMDAR autoantibodies often manifest SZ-like symptoms such as hallucinations, delusions and thought disorders (Vitaliani *et al*, 2005). Moreover, the presence of neuro-reactive antibodies have been demonstrated in various diseases involving the neuromuscular system such as systemic lupus erythematousus (SLE) (Hanly *et al*, 1994), Guillain-Barre syndrome (Yuki *et al*, 2012) and myasthenia gravis (Patrick *et al*, 1973). The presence of anti-Syncytin-1 antibody in SZ patients suggests the involvement of autoimmune mechanism in its pathogenesis as well. However, although expression of HERV-W has been previously demonstrated, there is no significant difference in the extent of its expression among SZ, BD and healthy subjects (Karlsson *et al*, 2001). Moreover, the presence of anti-Syncytin-1 antibody does not necessarily satisfy the classical and the revised postulates defining the autoimmunity (Rose *et al*, 1993). It is also noted that various factors known to induce HERV-W expression, such as infection with influenza and herpes viruses, are also recognized as risk factors for SZ (Li *et al*, 2014; Nellåker *et al*, 2006). These findings collectively suggest the link between SZ and autoimmunity although its role in SZ pathogenesis is not as straightforward as in other autoimmune diseases such as SLE.

Our present findings suggest the involvement of retrotransposons such as HERV-W in the pathophysiology of SZ (Karlsson *et al*, 2001; Perron *et al*, 2008; Perron *et al*, 2012). Previous reports demonstrated the expression of HERV sequences in brain (Yolken *et al*, 2000; Karlsson *et al*, 2001; Frank *et al*, 2005) and sequencing of the chromosome rearrangement site at the chromosomal break point revealed the involvement of HERVs (Feschotte *et al*, 2012; Weckselblatt *et al* ¸ 2015; Lee *et al*, 2015). Moreover, Ohnuki et al. reported the involvement of HERV-H activation in the differentiation from inducible multipotent stem (iPS) cells (Ohnuki *et al*, 2014). These findings suggest that the genetic rearrangement involves retrotransposons such as LINE and HERVs (Muotri *et al*, 2005; Evrony *et al*, 2012; Upton *et al*, 2015; Erwin *et al*, 2016). It is possible that aberrant or excessive activation of retrotransposons in neuronal cells and elicitation of host immune responses might be associated with neuropsychiatric disorders such as SZ. Gage et al. further postulated the active role of genetic mosaicism generated by such retrotransposons on the proper differentiation of neurons and construction of neuronal networks (Muotri *et al*, 2005; Erwin *et al*, 2016). A number of subsequent studies utilized a single cell genome sequencing technique and have revealed that up to 60% (Erwin *et al*, 2016), though still controversial (Evrony *et al*, 2016), of neuronal cells in healthy individuals undergo genetic rearrangement (Muotri *et al*, 2005; Evrony *et al*, 2012; Upton *et al*, 2015; Erwin *et al*, 2016). It is possible that the neuronal connectivity for the construction and maintenance of basic cognitive brain functions might have been impaired in SZ and BD brains. Interestingly, Bundo et al. reported that the frequency of L1 retrotransposition is higher in the prefrontal cortex of the post-mortem brains of SZ patients and that the L1-insertion was preferentially localized to synapse- and SZ-related genes (Bundo *et al*, 2014).

The activation of retrotransposons, including HERVs as well as LINE, might disorganize the directed genomic rearrangement of neuronal cells. This aberrant neuronal genetic rearrangement might deceive the decent formation of neuronal circuits by interfering with physiological synaptic connectivity, eventually leading to the development of neuropsychiatric disorders. Further molecular analyses are needed to clarify the relationship between the emergence of the anti-Syncytin-1 antibody and disease occurrence. Such studies should provide a novel approach for diagnosis and understanding of the pathogenesis of psychiatric diseases and could facilitate development of specific therapies involving the control of excessive retrotransposition using retroviral reverse transcriptase or integrase inhibitors.

## Materials and Methods

### Sera from patients and controls

Sera from patients with schizophrenia (SZ; 48 cases) and bipolar disorder (BD; 18 cases) were collected from hospitalized patients in Yahagigawa Hospital. Informed consent was obtained from each patient or their family after a thorough explanation of the purpose of the study. Sera from patients with multiple sclerosis (MS; 17 cases) and the control group (Cont; 79 cases) were similarly collected from Nagoya City University Hospital, Fuji Hospital and Tomita Hospital. Sera from patients under treatment with valproic acid (VPA; 7 cases) were collected after brain surgery in the Fuji Hospital. These sera were kept at 4 °C or frozen until use. Diagnoses of SZ and BD were made after the agreement of at least 2 psychiatric specialists using international criteria for disease classification, such as the DSM-5 (American Psychiatric Association, 2013) and ICD-10 (WHO, 1992). Patients with autoimmune diseases, cancer, conventional viral infections, or other psychiatric conditions were excluded from the study. This study was approved by the Medical Ethics Committee of each institution involved in the study (Approved number from Nagoya City University: No. 60-00-0886) and was conducted in accordance with the principles of the Helsinki Declaration.

### Synthetic DNA oligonucleotide and plasmids

The entire *syncytin-1* coding region of HERV-W (Mi *et al*, 2000; Lavialle *et al*, 2013) and the codon-substituted versions were synthesized enzymatically based on the nucleotide sequence, using the primers shown in Appendix Table S1. The 42 oligonucleotide fragments, each containing approximately 60 nucleotides with the overlap of about 20 nucleotide one other, were chemically synthesized to cover the entire *syncytin-1* ORF. Cloning of the codon-optimized gene was described previously (Nakamura *et al*, 2016). Briefly, these fragments were annealed, and the DNA fragment containing the *Syncytin-1* ORF was amplified by PCR. The amplified fragment was inserted into the bacterial expression plasmid pET-28a by ligation at the BamHI and EcoRI sites. This plasmid was thus named pET-Syncytin-1. The nucleotide sequence of this plasmid was confirmed through sequencing analysis.

### Antibodies

The following antibodies were commercially purchased from the indicated suppliers: anti-His antibody (Santa Cruz Biotechnology), anti-Syncytin-1 antibody (Santa Cruz Biotechnology) and horseradish-peroxidase (HRP)-conjugated antibodies against human or rabbit IgG (GE Healthcare).

### Purification of recombinant His-Syncytin-1 protein

The recombinant protein His-Syncytin-1 was expressed in *Escherichia coli* BL21-codonplus (DE3)-RIPL competent cells (Agilent Technologies), according to standard protocol. Cell pellets from the 1-L culture were resuspended in 20 mL of lysis buffer (50 mM Tris [pH 8.0], 5% glycerol, 1 mM DTT, protease inhibitor cocktail [Complete™ EDTA free; Roche]) and homogenized using a sonicator (Microson XL™; Micronix, Santa Ana, CA.). The lysate was centrifuged at 20,000 x g for 10 min. The supernatant was discarded, and the pellet was suspended in 40 ml of lysis buffer with 8 M urea. The lysate was subsequently centrifuged at 15,000 rpm for 10 min and the supernatant, containing the recombinant His-Syncytin-1 protein, was transferred to a new 50-mL tube. The His-Syncytin-1 protein was purified using HisTrap FF (QIAGEN) with a washing buffer (50 mM Tris [pH 8.0], 0.5 M NaCl, 20 mM imidazole, 6 M urea) and elution buffer (50 mM Tris [pH 8.0], 0.5 M NaCl, 200 mM imidazole, 6 M urea). To further purify the His-Syncytin-1 protein, the above semi-purified protein fraction was applied onto 10% SDS-PAGE and reverse stained (Fernandez-Patron *et al*, 2005), and the protein band detected at 55 kDa was excised, lysed in 2x SDS loading buffer, heated at 95 °C for 5 min, centrifuged, and recovered in the supernatant (Fig. 1).

### Western blotting

The purified His-Syncytin-1 protein was separated by 10% SDS-PAGE and transferred to a PVDF membrane (Immobilon-P™; Merck Millipore, Billerica, MA.). The membrane was immunoblotted with the patient’s serum (1:300 dilution in blocking buffer [5% skim milk and 0.1% Tween-20 in Tris-buffered saline (TBS)] or the indicated antibody (1:1000 dilutions in blocking buffer), and the bound anti-Syncytin-1 antibody was visualized with HRP-conjugated antibodies against human or rabbit IgG using a Super Signal West Pico Chemiluminescent™ substrate (Pierce).

### Spot synthesis of peptide microarrays for the epitope mapping

The cellulose membrane with 264 immobilized peptide spots was made by Biomine Solutions Inc, (Richmond, Canada). The spot synthesis was carried out according to the method from Gausepohl et al. using a fully automated protocol (Gausepoh *et al*, 2002). The peptide spots were coupled on the cellulose membrane. Each spot consists of a 12-amino acid (AA) peptide with a frameshift of 2 AA throughout the entire 538 AA based on the Syncytin-1 protein sequence. For epitope mapping using sera from the SZ patients, the membrane was first incubated with a blocking buffer (5% sucrose, 4% skim milk and 0.2% Tween-20 in TBS) at 25°C for 4 h and washed once in TBS-T (0.2% Tween-20 in TBS) for 5 min. The membrane was incubated overnight with a 1:300 dilution of patient’s serum or the control anti-Syncytin-1 antibody in blocking buffer, followed by three 20-min washes with TBS-T. The membrane was then incubated with a 1:4000 dilution of anti-human or rabbit IgG-HRP antibody in blocking buffer for 2 h at 25°C, and then washed with TBS-T thrice for 20 min. The reacted peptide on the microarray membrane was visualized by chemiluminescence as described above and scanned by ImageQuant LAS 4000 (GE healthcare) for determination of the photointensity of each spot. The spot intensity was then quantified using Image J software version 1.49 (https://imagej.nih.gov/ij/, Wayne Rasband, National Institutes of Health, USA). The relative intensity (%) of each spot was calculated using the sum of all spot intensities in each peptide microarray.

### Synthetic peptides for ELISA

The peptides used for ELISA were synthesized through a conventional method in Optimal Biotech Pte Ltd, Singapore. Their purity was assessed using high-performance liquid chromatography and mass spectrometry. Details of these synthetic peptides are described in Appendix Table S2.

### ELISA

To quantitatively determine the immunoreactivity of each serum sample, an Immobilizer™ Amino (Nunc) plate was coated with Syncytin-1 peptide #8 at 10 μg/mL in 100 mM sodium carbonate buffer [pH 9.6] at 25 °C for 2 h. Nonspecific binding sites were blocked by the blocking buffer (100 mM sodium carbonate buffer [pH 9.6], 10 mM ethanolamine-HCl) at 25 °C for 2 h. After washing, a 1:100 dilution of the patient’s serum in dilution buffer (1x PBS, 0.05% Tween-20, 0.1% NP-40, 3% skim milk) was incubated overnight at 4 °C. After washing, the sample was incubated with an HRP-conjugated secondary antibody against human IgG (GE Healthcare) (1:4,000 diluted) for 2 h at 25 °C. Subsequently, 100 μL of substrate buffer (0.1 mg/mL 3,3’,5,5’-Tetramethylbenzidine [TMB] substrate, 50 mM phosphate-citrate buffer [pH 5.0], 0.006% hydrogen peroxide) was added, and the plate was incubated at room temperature for 30 min in the dark, and the reaction was terminated using a stop buffer (1 M HCl). Absorbance was measured at 450 nm (reference 690 nm) using a Tecan Sunrise™ plate reader. In order to assure consistent washing, each wash was performed using 300 μl of wash buffer (1x PBS, 0.05% Tween-20, 0.1% NP-40). Five such washes were performed using a commercial plate washer (AMW-8; BioTec, Japan).

### Competitive ELISA

Competitive ELISA was carried out in order to confirm the specificity of the antibody epitope. Each serum (at 1:100 dilution) was incubated with 0.001–10 μg/mL concentrations of the competitor peptide for 2 h at 25 °C. The reaction was performed in the same manner as that of a standard ELISA assay. Subsequently, the sample was subjected to the same ELISA protocol.

### Statistical analysis

All statistical analyses were performed using R software version 3.3.2 (Stanford University Social Science Data and Software, Stanford, USA). The significance of the difference in the male to female ratio was analyzed using a χ^2^ test and the comparison of age distribution was performed using a Student t-test; p-values were calculated in both cases. The distribution of antibody titers detected by ELISA was analyzed using Wilcoxon’s rank sum test. A p-value of less than 0.05 was considered statistically significant, and all reported p-values were two-tailed. Receiver Operating Characteristic (ROC) curves were generated using the R statistical package and ROCR.

## Acknowledgements

We thank Ms. Sachiko Amano, Mr. Takaoki Gotoh, Ms. Harumi Tsujimura, Miss Juri Funasaka, Mr. Ryoki Ishihara and Mr. Ken Sasaki for their technical support and encouragement throughout this project. All the clinical samples were provided by the hospitals enrolled in this study. We also thank Ms. Sheriah L. M. de Paz for helpful comments during the preparation of the manuscript. This study has been financially supported by Grants-in-Aid for Research in Nagoya City University and the Junwakai Medical Corporation, Japan.

## Author contributions

YH and TO conducted most of the experiments and analysis and wrote the manuscript. YH, AK, HM, YN, TS and HK significantly contributed to the establishment of ELISA system for anti-Syncytin-1 antibody. TO, YH, KA, SN, KM, TA, NM, TF, YT and NI participated in this study for the diagnosis of patients enrolled in this study and contributed for collecting serum samples. All authors discussed the results and reviewed the manuscript.

## The Paper Explained

**PROBLEM:** Currently, there is no simple and objective measure for the diagnosis of schizophrenia.

**RESULT:** In this study, we have attempted to establish a novel diagnostic ELISA system for schizophrenia by examining the serum antibodies against Syncytin-1 protein encoded by HERV-W. We have identified the immune-reactive epitope against Syncytin-1 and established a simple and reproducible ELISA antibody screening assay system using the immune-dominant epitope peptide. Using this ELISA system, nearly 50% of patients with schizophrenia (24 out of 48 cases) exhibited seropositivity to Syncytin-1 whereas less than 4% exhibited positive in the control subjects (sensitivity 50.0%; specificity 96.2%).

**IMPACT:** Our results demonstrated not only the clinical feasibility for detecting the antibody against the Syncytin-1 protein, but also suggested the possible pathophysiological role of activation of human endogenous virus type W (HERV-W) in schizophrenia. Further molecular investigation of the involvement of retrotransposons such as HERV-W is warranted for understanding the disease occurrence and development of novel therapy.

